# An all-atom protein generative model

**DOI:** 10.1101/2023.05.24.542194

**Authors:** Alexander E. Chu, Lucy Cheng, Gina El Nesr, Minkai Xu, Po-Ssu Huang

## Abstract

Proteins mediate their functions through chemical interactions; modeling these interactions, which are typically through sidechains, is an important need in protein design. However, constructing an all-atom generative model requires an appropriate scheme for managing the jointly continuous and discrete nature of proteins encoded in the structure and sequence. We describe an all-atom diffusion model of protein structure, Protpardelle, which instantiates a “superposition” over the possible sidechain states, and collapses it to conduct reverse diffusion for sample generation. When combined with sequence design methods, our model is able to co-design all-atom protein structure and sequence. Generated proteins are of good quality under the typical quality, diversity, and novelty metrics, and sidechains reproduce the chemical features and behavior of natural proteins. Finally, we explore the potential of our model conduct all-atom protein design and scaffold functional motifs in a backbone- and rotamer-free way.

## 1. Introduction

Structure, or more specifically the set of spatial chemical interactions proteins make with themselves and other molecules, remains the best rationalizable connection between protein sequence and function. Therefore, the ability to design proteins for novel functions often involves some modeling of structure, implicitly or otherwise. In order to have the most control over the arrangement of structural elements, it is often beneficial to design proteins *de novo*, specifying fully the structure and sequence of a protein from scratch [1, 2].

Within a protein structure, sidechains are the primary functional effectors and define both the intrinsic properties of the protein and the types of interactions a protein can make. While it is well known that the sequence (and therefore the set of sidechain) determines the global fold of the protein, the interactions of sidechains with other sidechains can also influence local packing and structure, as well as distal allosteric effects. Sidechains also mediate most unique protein functions, ranging from protein-protein interactions to enzyme catalysis and metal coordination. Their precise placement in designed proteins is essential for downstream function. Combined with the backbone atoms, the ability to model the full atomic protein structure is important for advancing protein design.

However, modeling sidechains in protein design is difficult: to specify which sidechains need to be modeled, the sequence must be known, at which point the protein is already fully determined. Thus, current *de novo* design methods rarely model the sidechains at all when specifying the sequence or structure, but instead sample the backbone structure and sequence separately, and then build the sidechains afterwards [3–21]. Some other structure-oriented design methods allow the sequence and structure to interact during the design process, allowing for co-design and for each representation to influence the other [12, 22–24]. For some specific protein folds or classes such as antibodies, sequence and structure co-design has been explored, sometimes in an all-atom manner [25–29]. While these methods have led to remarkable advances in our ability to design protein structures and sequences with increasing control, robustness, and usefulness, most have not yet deeply explored the generation of protein structures conditioned explicitly on all-atom structure or sidechains.

Here we describe an all-atom protein generative model, Protpardelle, which co-designs the backbone, sequence, and sidechains of a protein together. Our method enables a key capacity in protein design: generating complete proteins with coherent sequences and all-atom structures. By co-designing the structure and sequence, the all-atom modeling approach allows the sequence to influence the backbone conformation through the sidechains, and vice versa. We accomplish this by developing a way to manage all of the sidechains at once during the generation process, which we dub “superposition” in reference to how quantum wavefunctions exist in multiple states before “collapsing” into a single state when observed. Model samples are of good quality, both in terms of consistency between structure and sequence as well as chemical fidelity of the sidechains. Preliminary exploration of design applications suggests that our model can be used to design new proteins in an all-atom context, as well as when conditioned on only the functional groups of protein sidechains. We also describe a performant backbone generative model as a special case of our model. Both models are computationally lightweight, which aids exploratory research; in service of this, we intend to make our code available at https://github.com/alexechu/protpardelle.

## 2 Method

### 2.1 A simplified ODE for protein structure modeling

Diffusion or score-based generative models [30–33] have emerged as a powerful framework for generating high-quality data samples in continuous domains, including protein structures [17, 18, 20, 21, 34–36]. They have shown promising results by utilizing an iterative generation mechanism that allows the model many opportunities to commit to and refine a sample. These methods are also highly amenable to conditioning, with several mechanisms to inject steering information and guidance; this is particularly relevant for protein design, since the final objective is almost always to produce a protein with some desired property [37–39]. Due to these attractive properties, we use the diffusion paradigm to construct our generative model.

To review the basic approach of diffusion-based generative models, we can define (1) forward and (2) reverse SDEs that connect an interesting distribution (e.g. the data distribution, *p*_0_(**x**)) to a tractable distribution (e.g. the isotropic Gaussian distribution, *p*_*T*_ (**x**)) [31]. The forward SDE reduces the signal to noise ratio until data is destroyed to whitened noise, and the reverse SDE recovers realistic data from random initial noise by progressively denoising noisy data.

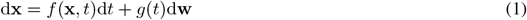

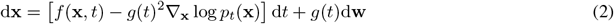

Here *g*(*t*) is a diffusion coefficient, **w** is the standard Wiener process, and *f*(**x**, *t*) is a drift term which is typically of the form *f*(*t*)**x** and describes a time-dependent scaling of the data. Different choices for the drift and diffusion coefficients give rise to the various variance-preserving and variance-exploding noise process formulations [31, 32]. Given a score model that computes or approximates the score, or gradient of the log density of data ∇_*x*_ log *p*(**x**), we can produce solutions to the reverse SDE, allowing us to generate data from noise. This score model is typically a neural network trained with denoising score matching, which we call *D*_*θ*_ [40, 41].

In place of the reverse SDE, we can also choose to integrate the equivalent probability flow ODE, which recovers the same marginal distributions [31]. One particular configuration of this ODE does not scale the data and uses the identity function for *σ*(*t*) = t so that the noise level increases at the same rate as time, or diffusion progress [42].

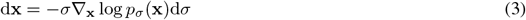

The marginals associated with this ODE are 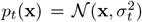, which can be interpreted as adding or removing Gaussian noise of constantly increasing scale during forward and reverse diffusion. In theory, this ODE should have more linear solution trajectories with reduced truncation error, and in practice we find it to be highly effective [42, 43]. Intuitively, integrating this ODE amounts to approximating the score with an estimate of the ground truth and then taking a small step Δ*σ* in this direction (Fig. 1A). (Since *σ* := *t* under this configuration, we will abuse our notation and use *σ, t*, and *σ*_*t*_ somewhat interchangeably to indicate the noise level or progress in the diffusion process. Occasionally we will use *t* − 1 to indicate the succeeding timestep during sampling even though our model is defined on continuous time.) We will use **x**_0_ to denote a sample from *p*_0_(**x**), i.e. unnoised data, and **x**_*t*_ to denote samples from the marginal distributions *p*_*t*_(**x**), i.e. noised data.

**Figure 1.**
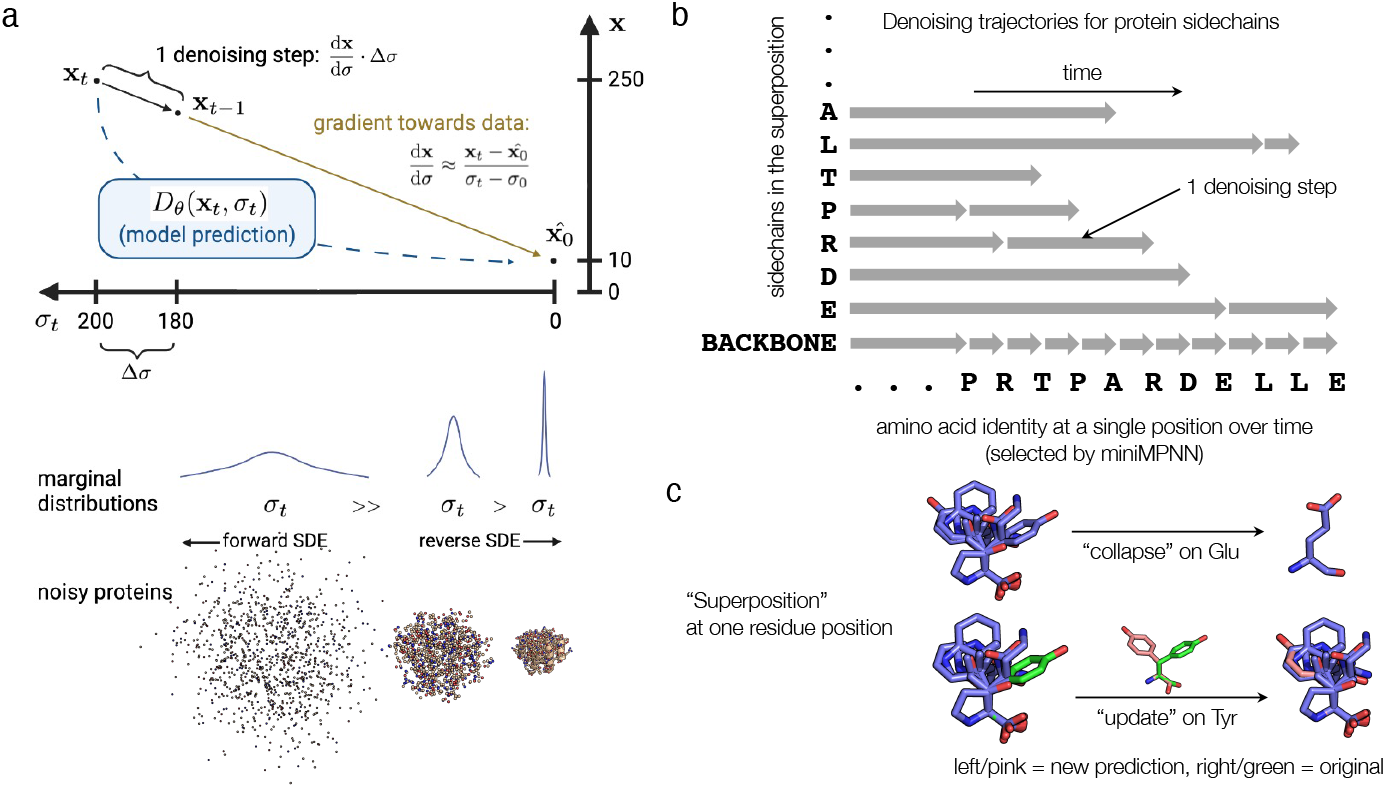
Superposition modeling approach and denoising scheme for Protpardelle. **(a)** The basic idea of denoising protein structures by integrating an ODE. Given noisy data x_*t*_, we can run the denoising network to predict the fully denoised data, x_0_. Given the quantities x_*t*_, x_0_, and the noise level *σ*_*t*_, we can estimate the score, or gradient which points in the direction of data. We can then take a denoising step (integrating the ODE) by choosing a step size Δ*σ* and computing an updated Δx on x_*t*_ which yields slightly denoised data x_*t*− 1_. We can repeat this many times to iteratively denoise our sample and produce protein samples. The noising process is defined by the marginal distributions, which noise protein structures by simply adding Gaussian noise to the atomic coordinates. The scale of these Gaussians increases linearly with time, thereby inducing mostly linear ODE solution trajectories. In our model, the forward noise process acts only on real proteins (with one sidechain per amino acid), whereas the reverse denoising process acts on the full “superposition” over all possible sidechains. **(b)** A visualization of the Protpardelle sampling routine for a single residue position. The vertical axis lists the structural elements being denoised (i.e. the atoms of the 20 sidechains in the superposition, plus the backbone atoms). The horizontal axis denotes progression in sampling time, with each amino acid denoting the amino acid predicted for this position at a given timestep. Note that this amino acid prediction can change from step to step. Briefly, at each timestep, we use the predicted amino acid to collapse the superposition and form a “real” yet noisy protein, predict denoised positions for each of the atoms in this protein, and then take a denoising step for selected atoms. The size of the denoising step for each atom or sidechain is determined by the last time that atom or sidechain took a denoising step. Each amino acid sidechain from the superposition is denoised only when it is selected by the sequence model. This means that the size of the denoising/integration step can vary depending on how frequently that amino acid is predicted. The backbone is denoised at every step since these atoms are common to all amino acids. For more details and the pseudocode of the sampling algorithm, see the Method section and Algorithm 1. **(c)** A visualization of the sidechain superposition idea and how it might be collapsed or updated at each denoising step. Sidechains for all 20 amino acids are modeled at once (here aligned on their N, CA, and C atoms). Given an amino acid type, we can collapse the superposition from all states to a single state, which yields a “valid” residue or protein. Alternatively, given an amino acid type and newly predicted coordinates for that sidechain, we can update the superposition with new information.

Protein structure data can be handled in many different ways; one simple and descriptive approach is to treat it as a point cloud in 3D space. To apply this ODE to noise protein structures, we would add Gaussian noise with variance 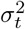 independently and identically to the x, y, and z coordinates of each atom in the structure (backbone or sidechain). As physical objects in 3D space, the atomic coordinates of protein structures obey the symmetries of rigid body rotation and translation (but not reflection). Thus our noising process should also consider the same symmetries. We note that the isotropic Gaussian distribution in *n* dimensions is symmetric and thus invariant to rotations, yielding an SO(n)-invariant density. Additionally, we always move the center of mass for all protein structures to the origin, which further ensures that the added noise is invariant to translation, yielding an SE(n)-invariant density [21]. This noise distribution induces an SE(n)-equivariant diffusion process [44], where the noisy protein structures **x**_*t*_ remain centered at the origin while rotating together with the original, true protein structure **x**_0_.

### 2.2 Sampling with an all-atom superposition

Training a score network to denoise protein backbone atoms (the N, CA, C, and O atoms) is straightforward, and we will also discuss our results with a backbone-only generative model. However, all-atom protein modeling presents an interesting challenge not only because of the dual continuous and discrete nature (structure and sequence) of proteins, but also because the discrete sequence directly defines which atoms are present in the 3D structure. For example, serine has five heavy atoms, N, CA, C, O, CB, and OG, whereas histidine has ten atoms, N, CA, C, O, CB, CG, CD2, ND1, CE1, and NE2. Five of these atoms (N, CA, C, O, CB) are common to both amino acids, but serine has an atom missing in histidine, and histidine has five atoms missing in serine. This creates a chicken-and-egg problem: for each position, we cannot know which sidechain atoms to build without knowing the amino acid identity, but if we know all the amino acid identities, then the protein is already specified. (Barring post-translational modifications and other biological processes, proteins are entirely determined by their sequence [45].)

Nearly all current generative modeling paradigms utilize deep neural networks which work with fixed-size input and output, with length and shape differences between unique data points handled by masking. For all-atom protein generation where both the protein structure and its sequence are unknown at the beginning of sampling, the estimated sequence evolves with time and therefore the structure consists of different atoms at each time step. Practically speaking, this means that not only does the data change due to the noise process, but the mask itself also changes with each diffusion timestep. This remains the case whether sidechains are represented as sets of atoms or as sequences of chi angles. This makes it difficult to define a diffusion process which transforms data smoothly with time, if the data is disappearing and reappearing at each step.

#### Algorithm 1

All-atom sampling

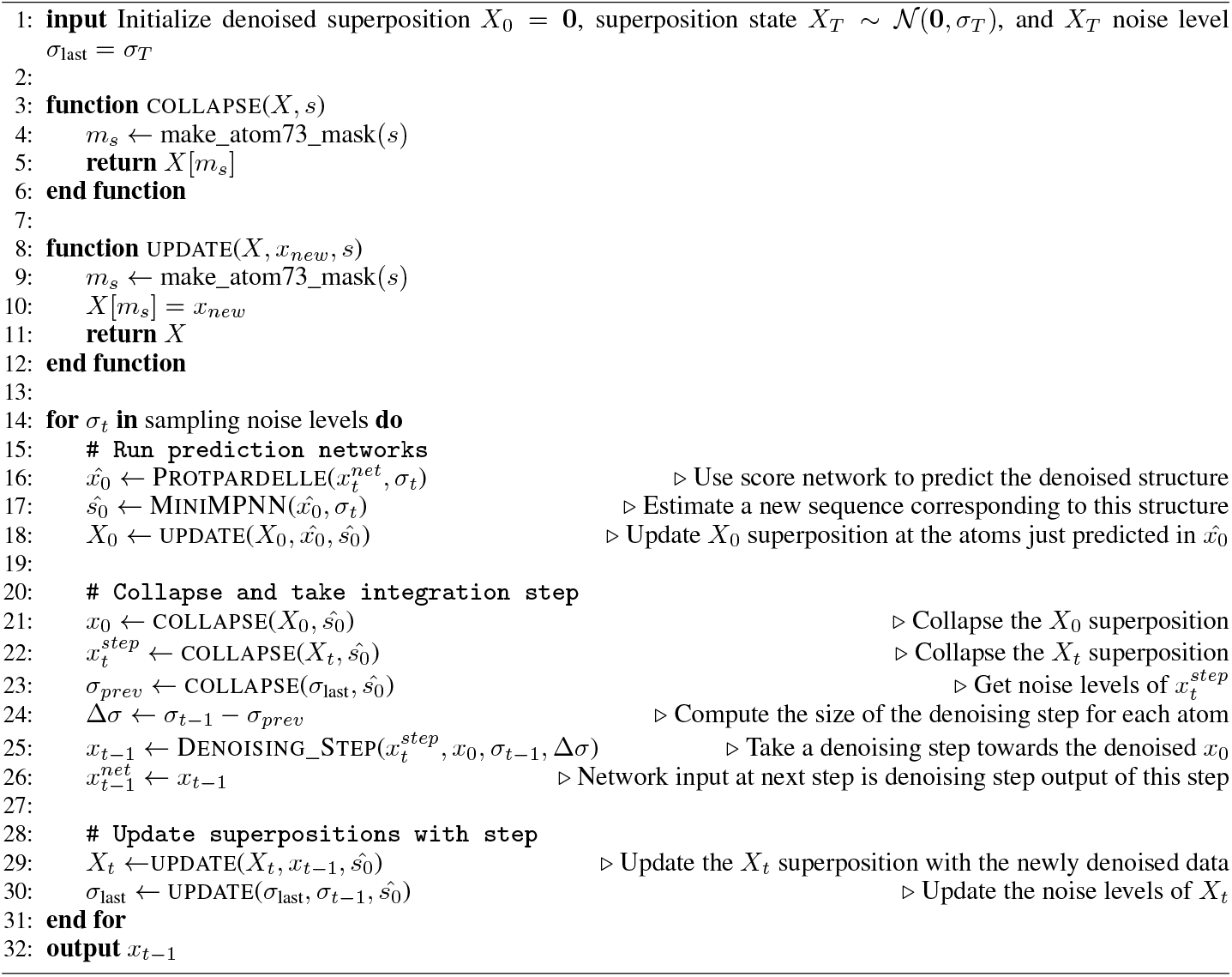

To address this challenge, we define our noising process to act on a “superposition” of protein structure states - that is, the protein backbone and the coordinates of each of the twenty possible sidechains at once. This is clearly an unrealistic model of protein structure, but allows us to handle the uncertainty associated with changes to the sequence in time. Given a sequence, we can “collapse” this superposition by selecting the sidechain states that correspond to this sequence to yield an all-atom protein structure. During structure generation, we maintain both an estimate of the fully denoised superposition state (*X*_0_) and the current noisy state (*X*_*t*_). At each denoising step, we collapse the *X*_*t*_ superposition to produce a single noisy protein structure *x*_*t*_ which we can use to predict the denoised data *x*_0_ with the score network. This predicted *x*_0_ can be used to update our *X*_0_ estimate and to predict a new sequence. Then, the actual denoising step occurs: we collapse X_*t*_ and X_0_ *using the new sequence* to get x_*t*_ and x_0_, and then we integrate the ODE using these two quantities (see Algorithms 1 and 2 for the full pseudocode). We use notations 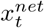 and 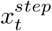 to distinguish the network input *x*_*t*_ and the denoising step input *x*_*t*_, respectively: note that the score network prediction and the denoising step are decoupled in our method. The output of the denoising step *x*_*t*−1_ at timestep *t* is exactly the 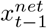 at the following timestep *t –* 1 trajectory; this arrangement is so that inputs to the network are always at the same noise level in the ODE solution

A key insight of our approach is that the integration step size can vary for different atoms, and the ODE discretization need not be identical for all atoms (Fig. 1B). In this approach, the backbone atoms (N, CA, C, and O) are denoised at every iteration of the algorithm, and the various sidechain atoms are denoised only when the sequence design model selects the corresponding amino acid for that position. Thus the sidechain atoms will typically see larger Δ*σ* integration steps than the backbone at any point in time. Mechanically, the superpositions are stored in an `“atom73”` representation which indexes the N, CA, C, CB, and O atoms and then each amino acid’s sidechains independently. The collapse and update functions are mask-based interactions with the `atom73` representation (Fig. 1C). To sample, we adapt a modified version of the stochastic sampling routine outlined in Karras et al. [42], which offers a high degree of flexibility to customize the sampling process. This routine uses the Euler method to integrate the ODE while injecting noise at each step (similar to Langevin dynamics or predictor-corrector methods) and accepts several tunable hyperparameters which we find can have a large effect on sample quality (see Appendix A.3 and Supp. Table S1 for more discussion).

### 2.3 Training the score network

Despite the fact that the model must manage all possible sidechain positions at once during the reverse process, the forward process does not require any superposition modeling at all, because the score is only predicted on the collapsed states during sampling. This means that we can noise real data to generate training examples, and the score network can be trained directly on these examples. This simplifies the training scheme significantly and enables many optimizations and experiments to occur cheaply at inference time.

Inputs to the model are samples from the marginal distributions 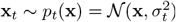, which can also be written as 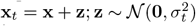 to highlight how noise is added to the data. We use only a single denoising score matching loss, with loss weighting:

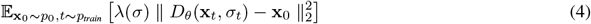

The score network *D*_*θ*_ is a simple U-ViT from computer vision which we augment with network preconditioning, a scaling scheme to streamline the training objective [42, 46]. In essence, inputs and outputs of the network are scaled and interpolated so that inputs are of consistent variance across training examples and noise levels. The loss weighting is determined by this preconditioning. Noise levels used to corrupt data were sampled from a log-normal distribution rather than the usual uniform distribution, which can be viewed as enriching the dataset at noise levels which are most critical for perceptual sample quality [42] (Appendix A.2). The loss function and network architecture are not equivariant to transformations in SE(3), but we suggest that in practice this property is not as important in generative modeling as it is in predictive modeling so long as the model is trained with the appropriate data augmentations (Appendix A.1-2). As such, inputs to the model have a random rotation and small random translation applied. We find benefits in the relative computational speed of non-equivariant networks. For more details on training, see Appendix A.2.

### 2.4 Sequence co-design

An all-atom generation approach also necessitates a way to estimate the correct sequence at each step of generation. In practice, any (fast) predictor of protein sequence given structure can fill this role. For this we used the ProteinMPNN graph neural network architecture which has been shown to capture an efficient locality-based inductive bias and produces strong sequence design results when used to parameterize an autoregressive model [47]. We adapted the architecture to produce a “mini-MPNN” model by removing the causal (autoregressive) mask which improves sampling time complexity significantly from O(N) to O(1) and augmenting the intermediate MLP layers with noise conditioning, allowing it to be trained on higher noise levels [48]. As input to the network we provide the denoised x_0_ structure and optionally the predicted sequence from the previous step, a strategy akin to self-conditioning [49]. With this approach, the sequence estimate becomes more and more accurate as the structure becomes better defined, so the “correct” sidechains are denoised more frequently as the trajectory progresses. We note that we do not define a diffusion process on the protein sequence; we only co-design the sequence with the structure.

It is possible for the sequence to influence the structure in two ways. One is the intended behavior where the positions of sidechains in space induce changes to the backbone to accommodate these sidechains. The second is for the model to infer the sequence from the atom mask and memorize the structure given this sequence, or “sequence leakage”. This creates a distribution shift issue during sampling where if the sequence is not plausible, the network is asked to denoise unusual inputs and struggles to produce valid structures. We remedy this by obscuring the atom mask by noising all 37 unique atom position inputs instead of only the atoms corresponding to the sequence. Later in the diffusion process as the structure information becomes clearer, the sequence predictions also improve and the problem recedes.

## 3 Results

### 3.1 Developing a method for all-atom generation

We first sought to establish the feasibility of the general ODE and denoising scheme by training the model on protein backbones only, i.e. generating the N, CA, C, and O atoms. Initially, we used a convolutional U-Net as the denoiser network and found helical structures to be easy to generate, but beta sheets to be more difficult to properly pair, likely due to their non-local nature. We found that changing the neural network to a U-ViT, removing the convolutional up- and down-sampling layers, and training with self-conditioning [46, 49] all contributed to reducing the frequency of helices and improving the quality of beta sheets. In general, we also found that network preconditioning and altering the training distribution over noise levels to focus on the most influential noise levels to be important strategies for improving sample quality [42]. One notable feature of diffusion models relative to other types of generative models is inference-time flexibility. We found that significant gains in sample quality can be obtained by tuning sampling hyperparameters, which is inexpensive (Table S1). We found that tuning the level of stochastic “churn” and a scale for the step size (similar to low-temperature sampling) during the sampling process has a major effect on sample quality.

With a baseline structure generation model in place, we next explored the capacity of the miniMPNN model to co-design the sequence during the structure diffusion. Following similar training procedures as for the original model, we found that miniMPNN was adequate as a structure-conditioned sequence predictor, achieving ∼38% sequence recovery on a validation set and a mean scRMSD of ∼8 in 1-5 sampling steps. We found that the base performance of the model as well as the many design steps (100s) used for structure diffusion led to lower quality sequences. To resolve this issue, we replaced the sequence prediction at the final step with a prediction from the full pre-trained ProteinMPNN [47]. This improved the sequence estimate and thus the self-consistency of the designed proteins. However the fast prediction of coherent sequences with miniMPNN was still needed during the sampling trajectory - full ProteinMPNN was too slow, but randomly selecting sequences also resulted in much lower quality samples due to the sequence leakage issue (see Methods 2.4).

With a basic approach in place for structure and sequence co-design, to enable all-atom protein generation, we needed to also diffuse the sidechains. The simplicity of our training scheme made this very straightforward: since our forward diffusion process is identical for each atom, we simply increased the number of atoms per residue. The model was able to generalize to this extension easily. However, when examining more closely some of the early failure modes for sidechain generation, we noticed that most of the sidechains were generated to a reasonable degree of chemical fidelity, but a small percentage of sidechains showed some divergence during sampling, producing bond lengths that were wrong by several angstroms. We hypothesized this was because the sidechains that were infrequently selected during sampling had either too few denoising steps overall, or perhaps a single denoising step that was too large. We did not find a significant association between the largest denoising step and sidechain divergence, but we did note that chemically invalid sidechains seemed to be sampled less frequently on average. To resolve this issue, we annealed the sequence resampling rate with time to increase the rate at which likely sidechains were selected; this improved the overall quality of the generated sidechains and structures. Additionally, we found that running a second sampling stage conditioned on both the backbone and sequence (i.e., essentially conducting rotamer packing) entirely removed any faulty sidechains. This second stage added very little sampling time since it did not involve running miniMPNN or ProteinMPNN. Together, these modifications and improvements allowed us to consistently generate high-quality all-atom protein structures.

### 3.2 Evaluating the generative model

We evaluated our model on three main properties which are relevant for generative models, all related to sampling: the quality (broadly defined), diversity, and novelty of model samples. Intuitively, we desire our model to generate “good” samples that exhibit structural diversity and can generalize beyond the training set. The plausibility (or designability) of sampled proteins is clearly important because we want our designs to fold successfully in solution, and can be evaluated using self-consistency metrics, which predict the structure of a designed sequence for a protein and assess the agreement between the predicted structure and sampled structure. The agreement is typically scored with either the RMSD metric or the TM-score metric calculated on the alpha-carbon atoms (referred to as scRMSD and scTM), and these metrics have been suggested to correlate with experimental success [13, 18, 21]. We compute these scores on both the backbone and all-atom models using ESMFold as a structure prediction oracle and ProteinMPNN as a sequence design model [47, 50]. We can further assess the chemical quality of model generations by measuring quantities such as bond lengths, bond angles, and dihedral angles, and assessing whether they align with real proteins. In this vein we also compute the mean bond length RMSE metric, which measures the mean deviation of each bond length from an ideal value, and which we find correlates with the other angle-based statistics.

The diversity of model samples is an important quality not only because we want to avoid artifacts of generative modeling (e.g. mode collapse, where models optimize their training objectives by producing a limited range of high-quality samples), but also because we want to draw deeply on protein structural space to produce solutions for different design problems. Diversity can be examined by clustering samples and counting them or measuring the maximum TM-score between clusters, but can be a difficult property to assess in general since it is closely tied with the number of samples drawn. We chose to cluster our samples and count the number of clusters, following the previous approach of Yim et al. [21]. We also computed the secondary structure content of samples with DSSP, measuring whether samples cover a broad range of alpha and beta-type structures [51]. Finally, to assess whether our model is able to generalize beyond the dataset (i.e. to evaluate novelty), we measured the TM-score of each sample against its nearest neighbor in the dataset. This can be done efficiently with FoldSeek, an algorithm which enables fast structure alignments at scale [52]. This nnTM metric indicates whether a sample is memorized or reproduced from the dataset, and describes the model’s ability to produce entirely novel proteins, relative to its training set.

The backbone model, when combined with ProteinMPNN as the sequence design module, achieves strong performance on these properties. Our sampling-time tuning and optimization allows us to unconditionally generate proteins of length up to 300 with comparable designability to current methods, at a much lower computational cost (Fig. 2A-C). Our success rate under both the scRMSD and scTM metrics is *≥* 90% for most of this length range, dropping to ∼70% at the top end, while generating samples up to one to two orders of magnitude faster than other current methods (∼0.002 seconds/residue on NVIDIA A40, compared to 0.04-1.5 seconds/residue on A100/A4000 [18, 21]). At lengths above 300, the model is still able to generate good proteins, but the success rate drops off rapidly. This could be for many reasons, including that there are much fewer proteins of this length in the dataset (Supp. Fig. S1B), that more ProteinMPNN sequences are needed as the error rate compounds with the sequence length for autoregressive models, or simply that the problem is harder. Samples are also quite diverse, covering a broad range of alpha-beta content proportions (Fig. 2E). As expected, the number of unique structure clusters grows with the number of samples, but the ratio of these two numbers decreases with the number of samples (Fig. 2D). With respect to novelty, given enough samples, the backbone model is able to on occasion generate proteins that belong to unique folds compared to those found in the training set (with TM < 0.5) (Fig. 2F). However, most nnTM values fall between 0.6 and 0.8, indicating that most samples share a common fold with a dataset member, but never just reproduce training examples. We also briefly explored inpainting and scaffolding motifs at a backbone-only level, and find that we are able to generate reasonable inpainting designs (Supp. Fig. S4).

**Figure 2.**
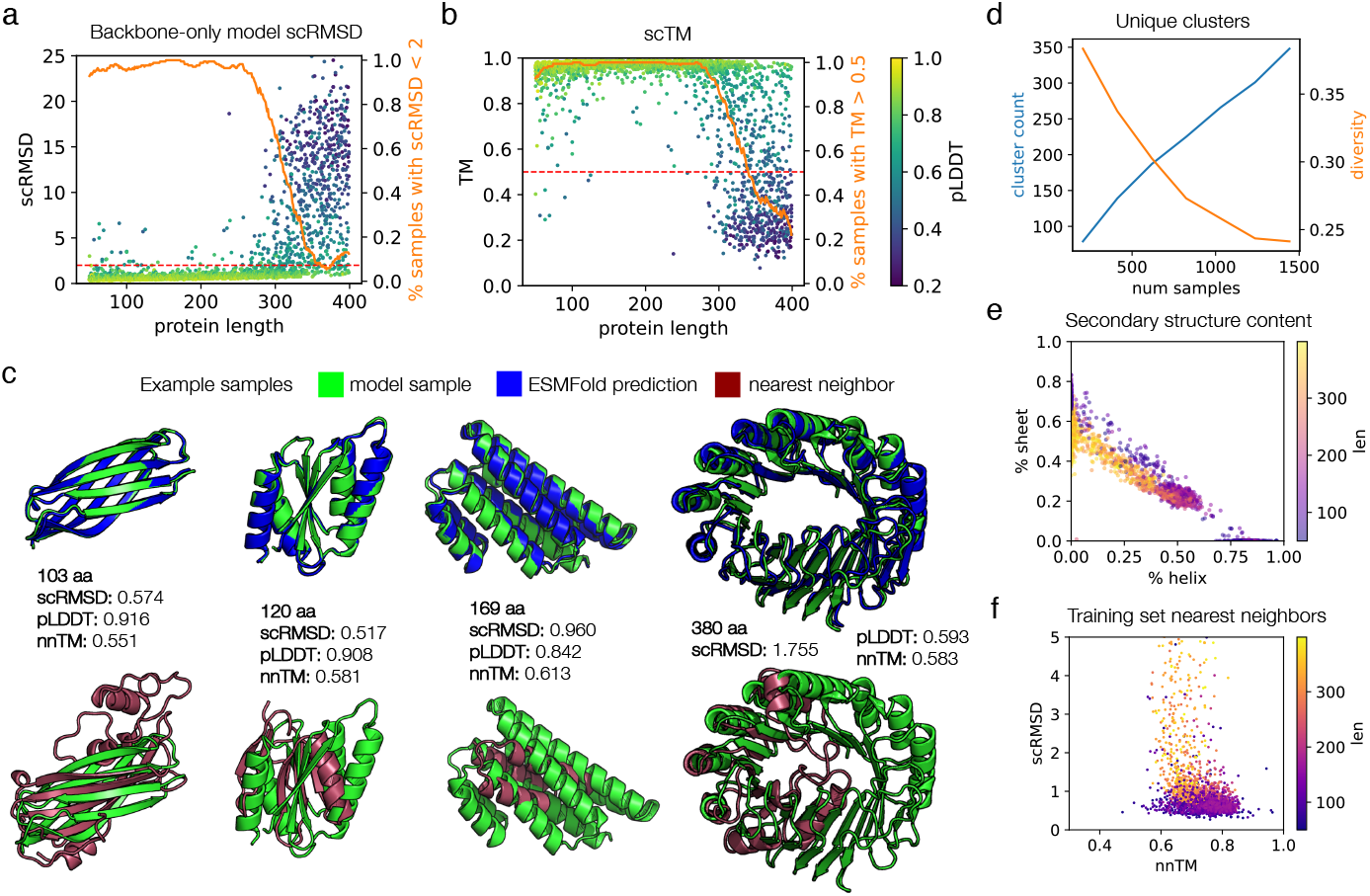
Evaluation of proteins sampled from the backbone-only model. **(a)** Self-consistency performance. We show the RMSD = 2 threshold (dashed line) and the proportion of samples passing this threshold, smoothed with a sliding window of size 11 (solid line). Eight backbones were sampled for each length from 50 to 400. For each backbone, the best of 8 ProteinMPNN-designed sequences is selected and ESMFold is used for all structure predictions. **(b)** The same samples and ESMFold predictions as in **(a)**, but using the scTM metric. TM is computed using the same alignment as for RMSD. Dashed line indicates TM = 0.5; solid line indicates proportion of samples with TM > 0.5, smoothed with sliding window of size 11. **(c)** Example high-quality, novel backbone model samples in green, shown aligned to the ESMFold prediction (blue) and the nearest neighbor in the dataset (red). The lengths, scRMSD, pLDDT, and nnTM metrics for each sample are also shown. **(d)** Number of structure clusters per samples drawn (left axis), and ratio of number of clusters to number of samples (right axis). Samples drawn uniformly over each length from 50 to 256 are used for this plot; the first 1, 2, 3, … samples for each length are used (so the number of samples is 206, 412, 618, … and so on, and there is always the same number of proteins for each length). **(e)** Secondary structure content of samples, computed by DSSP. **(f)** Nearest neighbor distances for model samples with scRMSD < 5. The nnTM is the TM score against the dataset member with the highest TM score to the sample, extracted with Foldseek.

We assessed the all-atom model on the same metrics to evaluate its ability for unconditional protein generation. Sampling is fairly robust at lengths up to 150, with a success rate of ∼60% on proteins in this range when assessed by scRMSD (Fig. 3A,B). We are able to retain high sampling speeds, though the model is approximately ten-fold slower than the backbone-only model, due to the need to run miniMPNN (Table S1). Visually, these samples cover diverse types of protein folds and are rich in secondary structure content (Fig. 3C). Above this length, samples do not become dramatically worse, but instead the average scRMSD shifts upward by 1-2 angstroms, leading to many samples which are “okay” but do not meet the scRMSD <2 quality threshold. If we examine the less stringent scTM score, the all-atom model is able to generate proteins for which the sequence adopts the correct fold (TM > 0.5) over 80% of the time throughout the full range of lengths tested. We observe that the scRMSD metric is able to distinguish between the more consistent model performance at <150 residues and the weaker performance above this length; whereas under the scTM metric the model seems to perform equally well across the full length range. In general, the pLDDT appears to correlate well with scRMSD for both models (Fig. 2A, 3A). Although the folding landscape as evaluated by these metrics suggests a degree of sequence-structural specificity, we cannot rule out potential overfitting of the predictive models [53]. Compared to the backbone model, the all-atom model appears less robust at all lengths, even when comparing only one ProteinMPNN sequence designed for each backbone-only sample (Supp. Fig. S1A). This manifests in both the scRMSD and pLDDT metrics. This suggests that modeling protein backbones becomes more difficult for the all-atom model, perhaps because of the additional complexity of modeling the sidechain atoms. Despite the fact that we do not explicitly provide sequence to the model, it is possible that sample quality and/or diversity may be affected by sequence leakage. (Methods 2.4).

**Figure 3.**
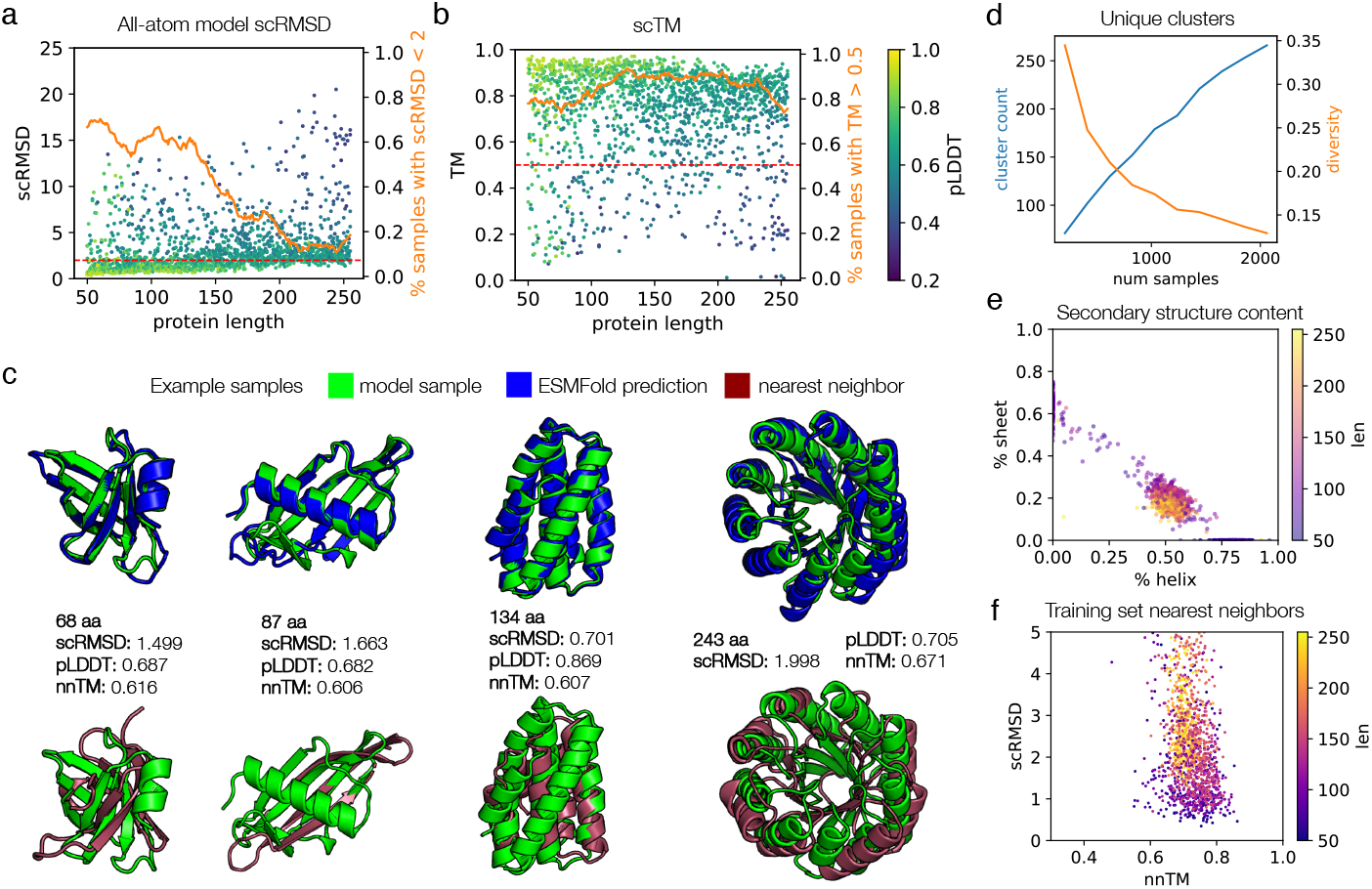
Evaluation of proteins sampled from the all-atom model. **(a)** Self-consistency performance computed as in Fig. 1, but for the all-atom model. Eight proteins were sampled for each length from 50 to 256. Each protein’s sequence is used for ESMFold, i.e. only one sequence is predicted for each sample, rather than the 8 sequences per sample that were predicted for the backbone-only model samples. The success proportion line is smoothed with a sliding window of 21. **(b)** The same samples and ESMFold predictions as in (a), but using the scTM metric. **(c)** Example high-quality, novel all-atom model samples. **(d)**Number of structure clusters per samples drawn (left axis), and ratio of number of clusters to number of samples (right axis). Samples are drawn uniformly over each length from 50 to 256, as in Fig. 2. **(e)** Secondary structure content of samples, computed by DSSP. **(f)** Nearest neighbor distances for model samples with scRMSD < 5. The nnTM is the TM score against the dataset member with the highest TM score to the sample, extracted with FoldSeek.

Model samples also exhibit diversity and novelty. When we draw more and more samples uniformly over the length range, we are able to continue to produce new structural clusters, although the ratio drops off at larger numbers of samples (Fig. 3D). Secondary structure content is also diverse across both alpha helix and beta sheet structures, and we note that we are able to generate proteins of all compositions (Fig. 3C,E). However, we notice a stronger relative preference for proteins with roughly 50% helix and 20% sheet content. This could be due to the fact that we only train on cropped proteins up to length 256 and lose the diversity at higher lengths (compare Fig. 2E and 3E), overfitting to the dataset at certain lengths where data is more scarce (Supp. Fig. S1B), or sequence leakage (Methods 2.4), and is a direction for further investigation. When searching for training set nearest neighbors, we see the same overall pattern as observed for the backbone model, with few memorized samples and few completely novel samples (Fig. 3F).

The previously discussed metrics can be computed solely on a structure (i.e. backbone) and the corresponding sequence. Since our model generates the atoms of the sidechains independently and does not enforce idealized bond geometries, we further evaluated the chemical quality of the samples. When compared to the ground truth training data, we find that samples from the model generally follow the same distribution with the same modes for bond lengths and bond angles, but with greater variance, as is often the case with free-atom generation methods (Fig. 4A,B). We conduct these analyses without relaxing model samples or the dataset under an energy function such as Rosetta, since the noising data augmentation in diffusion destroys this information anyway [54–56] (Appendix A.2). When examining the chi angles, the model samples are able to capture the two main modes of the natural distribution (Fig. 4C). The model distribution is more smoothed at lower values, missing one of the smaller modes and showing greater density than natural proteins in some regions. When we visually examine the generated structures, they appear plausible, exhibiting convincing packing and sidechain rotamers, and in some cases reproduce specific sidechain interactions seen in natural proteins, such as salt bridges between charged surface residues, helix capping, and some hydrogen bonding interactions (Fig. 4D).

**Figure 4.**
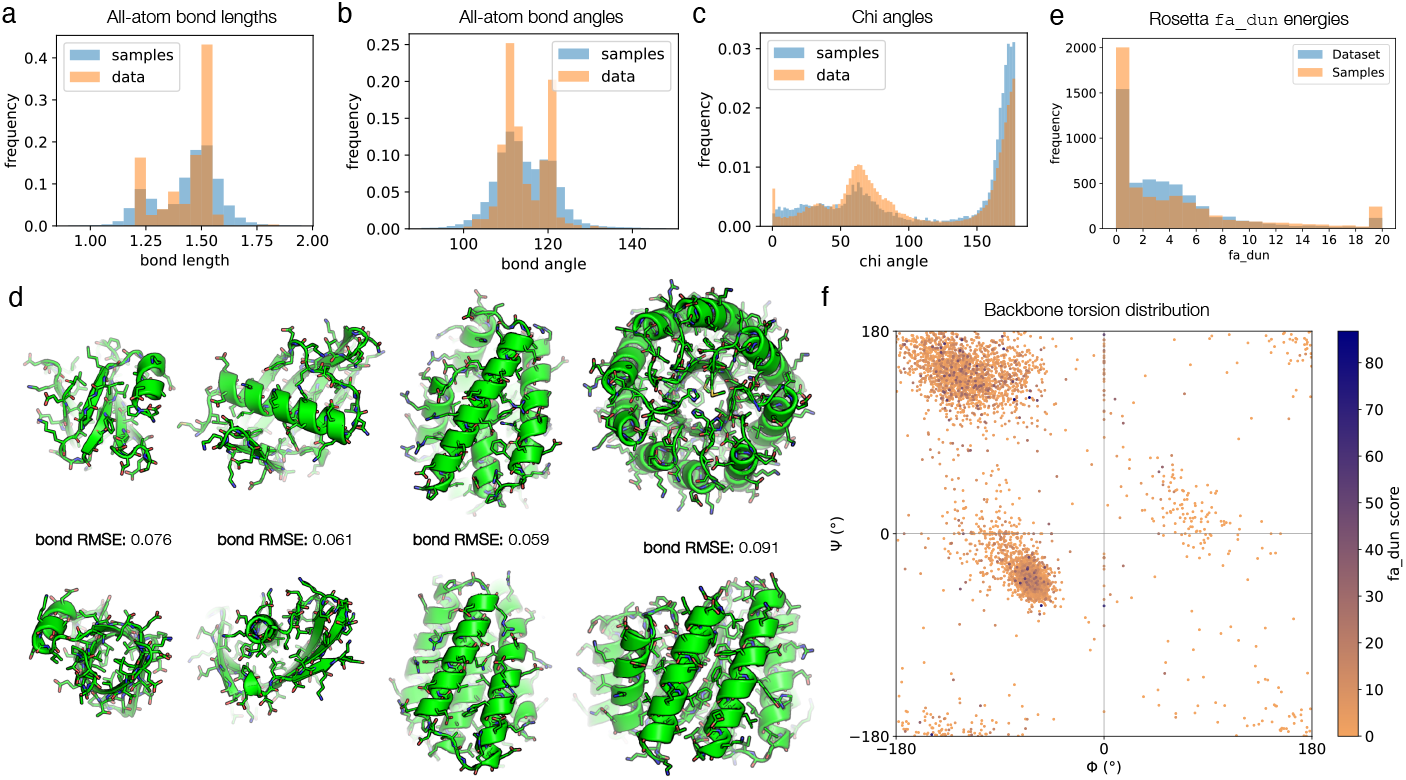
Analysis of generated all-atom structures, including sidechains. **(a-c)** Comparison of distributions of (a) bond lengths, (b) bond angles, and (c) chi angles for training data and model samples. Quantities for real data are computed from 100 random proteins from the training set; quantities for model samples are computed from 1 sample for each even-numbered protein length from 50 to 256. **(d)** Detailed views of all-atom raw model output with sidechains built. These are the same proteins as those shown in Fig. 3, only with sidechains rendered. The bond length RMSE is shown, which is computed by averaging the RMSE between each individual bond length and an idealized bond length in angstroms. For comparison, unrelaxed structures of natural proteins typically have an average bond length RMSE of 0.01-0.02. **(e)** Distribution of `fa_dun` energies for model samples and natural proteins. Statistics are computed from 5000 residues chosen at random (without regard to individual proteins) each from the dataset and the set of model samples. The `fa_dun` energy is computed from the probability of a rotamer given the backbone torsions and a potential term for deviation from an ideal chi angle value. **(f)** Visualizing the model samples data from (e) on a Ramachandran plot. Each point is a pair of residue backbone torsions, colored by the `fa_dun` Rosetta energy.

We also wanted to examine some of these properties statistically and explore whether the model learns to reproduce the backbone-dependent rotamer distributions observed in natural proteins and recorded in the Dunbrack rotamer libraries [57]. For each residue in a set of model samples, we computed the backbone phi-psi torsion angles and the `fa_dun` Rosetta energy term (a score derived from the probabilities of the rotamers and harmonic potentials for the chi angles, given the phi-psi backbone torsions). Without relax, most residues score within a tolerable range for the `fa_dun` energy, and closely follow the distribution of (unrelaxed) natural protein structures (Fig. 4E). Overall, the model samples obey proper chirality rules and exhibit backbone torsion distributions similar to native proteins (Fig. 4F). Outliers in `fa_dun` energy do not seem to correlate with any particular backbone torsion bin, suggesting that the model can generate sidechains for all forms of secondary structure well (Fig. 4F). We note that the `fa_dun` energy term can be noisy; in some cases the score is very high, but this is also observed in natural proteins (Fig. 4E).

### 3.3 Sidechain-conditional protein design

Our fundamental motivation was to develop a method for protein design that factors all-atom information into and throughout the entire design process, and to move towards design methods that allow for conditioning on arbitrary portions of protein structure, such as functional chemical groups, in a backbone- and rotamer-independent way. To this end, we explored whether the model has potential for designing new proteins in an all-atom manner. We trained a preliminary, crop-conditional model by providing it with randomly selected residues. These crops included contiguous spans of residues and discontiguous yet proximal residues; for some of these examples, the backbone was masked so that only the sidechain was provided to the model.

To test our ability to design new protein-protein interactions, we explored scaffolding a protein-binding motif in an all-atom manner, providing as input the backbone, sidechains, and sequence of only the motif. For this, we selected a *de novo* monobody designed to bind TGF-*β*1, a cytokine that activates a receptor kinase with multiple downstream signaling targets (PDB: 4KV5). Since this protein is designed *de novo* by the Sculptor algorithm [19] and and does not have a crystal structure in the PDB, it is guaranteed to not have been seen by the model during training. We extracted a loop and its sidechains from the binding motif of the monobody as conditioning input to the model and generated samples using Protpardelle. The model successfully generated protein structures that scaffold this motif and exhibit folds different from the original monobody (Fig. 5A).

**Figure 5.**
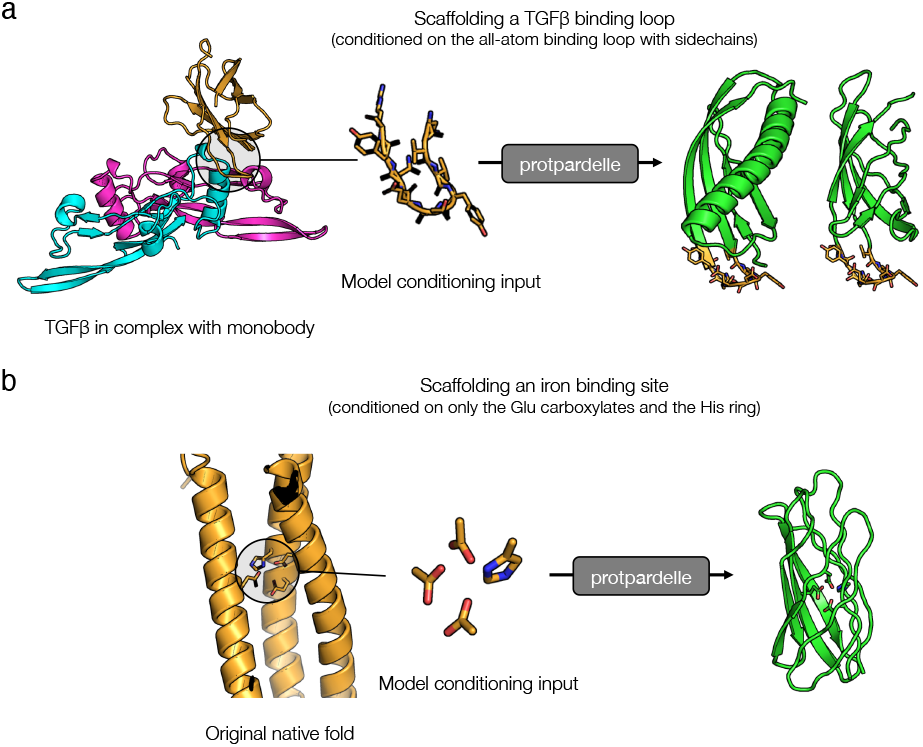
Towards all-atom protein design. Potential applications of our model for new approaches to protein design. The conditioning portion is shown in gold on the model sample (green indicates the model-generated portion). These designs are generated with an initial crop-conditional model and reconstruction guidance. **(a)** An example design generated by scaffolding a TGF-*β*1 binding loop including its sidechains. The original binder design (in gold) is a *de novo* designed monobody and thus is guaranteed not to be in the training set. The pink and cyan chains are TGF-*β*1 (PDB: 4KV5). **(b)** An example design generated by scaffolding only the functional groups of iron-binding Glu and His residues. The model is given only the atoms after the last chi angle: (CG, CD, OE1, OE2) for the Glu residues and (CB, CG, CD2, CE1, ND1, NE2) for the His residues. The original native fold is chain A of 1BCF.

A method to generate protein structures in an all-atom manner also enables new ways to generate proteins conditioned on functional motifs. Most current machine learning-based methods rely on models to first infer backbone conformations that seem mutually probable with the binding targets, and then to design the sequence and model the sidechains. We wanted to explore whether it is possible to generate complete proteins conditioned directly on the chemical groups which mediate the interaction. A metal-binding interaction is primarily dependent on the ligand interactions formed by the polarizable groups on the surrounding sidechains. We extracted only these polarizable groups for a single metal from a natural diiron-binding protein, cytochrome b1 (PDB: 1BCF); these groups included the carboxylates on three glutamate residues and the imidazole ring from the histidine residue. With only the atoms of these specific groups as conditioning input to the model, the model was able to design a protein scaffold that hosts these groups, together with the remaining sidechain atoms and rotamers needed (Fig. 5B). Some designed folds also differ from the original native fold. This suggests that our model could be a new approach to designing functional proteins in a backbone- and rotamer-free manner.

In these cases, we are able to obtain solutions resembling plausible proteins and indicate that this method could be a promising new approach for protein design. However, fidelity to conditioning, sampling success rate, and overall quality of these conditional generations remains to be improved. While standard replacement guidance was sufficient for conditioning tasks with strong input information, such as loop inpainting, we found that for tasks such as motif scaffolding with much less input information, the model was prone to ignoring the conditioning during generation. For these tasks, reconstruction guidance was needed to improve the coherence of generated designs with the provided conditioning information [39]. Many protein design tasks can be posed as some form of inpainting; training crop-conditional models has been a successful strategy for inpainting and scaffolding continuous and discontinuous functional sites elsewhere [11, 18, 24]. We expect this to hold for Protpardelle, as well as the many other tactics for applying conditioning in score-based generative models.

## 4 Discussion

Our results describe a number of contributions that we hope will advance the field of protein design. We have applied a new diffusion process and network architecture, both originally developed in computer vision, to protein structures in a way that allows for high-quality sampling of proteins. We describe a way to alter this process during sampling with a sidechain “superposition” so that a model trained only on standard protein structures is able to denoise sidechains for arbitrary sequences that any sequence design model might choose. This enables all-atom protein generation in a way where the model is able to reason about the sidechains jointly with the backbone, since both are noised and denoised together. Finally, we discuss potential extensions of this model to enable new ways to design proteins without dependence on modeling structural interactions implicitly through the backbone.

We stress that while our model is capable of co-designing sequence and structure, it remains a structure-primary generative model that produces estimates of the sequence during its sampling trajectory. It does not define any noising process on the sequence; nor is it a joint model in the sense that we are able to marginalize and condition in some way to produce solutions to the sub-tasks of structure and sequence generation and forward and inverse folding. However, structure-primary approaches have shown ever increasing capabilities to generate proteins with novel functions [18]. We hope that as sequence co-design and all-atom modeling become integrated as we have shown here, new ways to solve difficult protein design goals can be found.

While we have primarily explored applications for generating new proteins and designing proteins conditioned on explicit sidechain interactions, we note that our model automatically incorporates new ways of solving old protein design problems. In particular, we note that rotamer packing is an easy and efficient special conditioning case of this model. This capacity is demonstrated briefly in the second-stage design of our sampling process, where the sequence and backbone are fixed but the sidechains are re-sampled. We believe this to also be an interesting direction for future methods development.

## 5 Author Contributions and Acknowledgements

Conceived the project and designed the research: A.E.C. and P.-S.H. Developed the method and trained the models: A.E.C. Evaluation experiments: L.C. and G.E.N. Generated designs: A.E.C. and G.E.N. Wrote the manuscript: A.E.C. and P.-S.H. Contributed figures: A.E.C., L.C., and G.E.N. Editing and formatting: G.E.N., M.X., and A.E.C. Guidance and feedback: all authors.

We would like to acknowledge Phil Wang for generous sharing of many useful Pytorch modules, Kilian Cavalotti for help with the Sherlock computing cluster, Christian Choe for providing monobody models, and Daniel Richman, Jiaming Song, Simon Kohl, Russ Bates, Rob Fergus, Jonas Adler, and Sander Dieleman for helpful discussions. A.E.C and G.E.N. are supported by NSF Graduate Fellowships, and M.X. is supported by a Sequoia Capital Stanford Graduate Fellowship. A.E.C is additionally supported by Merck SEEDs Program. P.-S.H. is supported by NIH (R01GM147893), American Cancer Society (ACS 134055-IRG-218), BASF CARA project, and Discovery Innovation Fund. We would also like to thank an old roommate whose comment on a PyMOL session “whoa, are we designing pastas?” inspired the model’s name.

## A. Appendix

### A.1 Network architectures

The structure network has a U-ViT architecture [46] with a hidden dimension of 256 and is composed of 6 residual noise-conditional transformer layers. The self-attention layers contain 8 attention heads of dimension 32, and the feedforward layers are 3-layer MLPs with an intermediate hidden dimension of 1024. Typically the U-ViT architecture is trained with convolutional down- and up-sampling layers, though we remove them; we find that adding these layers improves alpha helix generation but slightly diminishes the quality of beta sheets. We use a patch size of 1 x n_*atoms*_ so that each residue is its own patch. We apply preconditioning from Karras et al. which stabilizes network training [42]. Inputs to the model are the 37 × 3 input noisy atom coordinates for each residue, the 37 × 3 self-conditioning atom coordinates, the noise level, and the sequence mask (but not the atom mask). We use absolute position encodings but also experimented with relative and rotary embeddings and find them to work well. We found training with mixed precision and weight decay to be harmful to performance. For the crop-conditional model, we provide additional 37 × 3 crop-conditioning coordinates. We explored providing the miniMPNN sequence prediction as input, but this had a minimal effect. Both the backbone-only model and the all-atom model are constructed with the same general configuration described here, with exception of the embedding layer which has a larger dimension in the all-atom model to account for the additional input atoms. We did not deeply explore the effect of tuning the network size hyperparameters.

The model is not fully SE(3)-equivariant, but this largely does not affect the interesting use cases of the model. Providing structural information through an attention bias presents an invariant way of conditioning, and providing crop conditioning to the model defines the rotational frame for the sampling process. Finally, since our diffusion processes are based on SE(3)-invariant densities, we note that converting the model to a fully SE(3)-equivariant one is only a matter of replacing the U-ViT module with an equivariant module which operates on 3D coordinates.

To inject noise conditioning information, we apply a noise-dependent affine transformation (scale and shift) to network activations [48]. These are applied to the inputs of attention layers after normalizing, and to the intermediate representations of MLPs in feedforward layers, in miniMPNN, and in the distogram projections.

The miniMPNN model uses the same overall configuration as ProteinMPNN. We made two modifications: we removed the autogressive mask and added noise conditioning to the MLP blocks, and then trained it from scratch. We skewed the noise applied to the coordinates heavily towards 0 as we found that the model does not train effectively if it is frequently shown very noisy structures. While we currently use ProteinMPNN for the final step, we believe a mini-MPNN-only approach to be feasible as well; strong sequence design results have been obtained with non-autoregressive (albeit not noise-conditional) ProteinMPNN elsewhere [58].

### A.2 Training details

We trained on the CATH S40 dataset which extracts domains from the PDB and removes redundant domains with *≥* 40% sequence identity [59]. We split the dataset into train, validation, and test sets using the splits in [60] and used the validation set to compute denoising loss metrics during training and evaluate the miniMPNN model. The backbone model is trained on crops of length up to 512 and the all-atom model is trained on crops of length of to 256. We do not relax the dataset under an energy function since the data augmentation associated with training diffusion models largely erases this information, although this can be explored in future work.

The model is trained with the Adam optimizer [61] with learning rate 1e-4 with a batch size of 32. We used linear warmup for the learning rate for 1000 steps followed by cosine decay. On a single NVIDIA A40 GPU, this takes approximate 2 days for 1 million iterations for the backbone model, trained on proteins cropped to length 512, and 3 days for 2 million iterations for the all-atom model, trained on proteins cropped to length 256. Training for the structure model was completed in 2-3 days on a single NVIDIA A40 GPU. We note that this relatively low computational commitment indicates it may yet be possible to obtain significant performance improvements by scaling the model and dataset.

We also used a log-normal distribution *p*_*train*_(*σ*) over the noise levels which we find important for successful training [42]. Noise levels applied to the data during training are sampled from a log-normal distribution over the noise levels, with *μ* = −1 and σ = 1.5. In detail, we draw a sample from 𝒩(− 1, 1.5^2^), exponentiate it, and scale it by the standard deviation of the data to determine the scale of the Gaussian noise to add. (For the backbone model, we use 𝒩(− 1.2, 1.2^2^).) A unique noise level is sampled for each minibatch element. Data augmentation is applied by centering the protein structure at the mean alpha-carbon coordinates, sampling and applying a rotation matrix uniformly at random, and sampling and applying a random translation (3-vector) from the standard normal distribution.

Additionally denoising the sidechains might have a similar effect as diffusion modeling with high-resolution images; since there more atoms, more noise might be needed to destroy the same amount of information [62]. We did not explore customized noise schedules, such as distinct or correlated noise schedules for backbone and sidechains, beyond a simple scaling of the noise scale for the sidechains that did not appear to be beneficial. Defining this relationship more clearly might enable tuning the trade-off between backbone and sidechain quality and the degree of interaction between the two during sampling.

### A.3 Sampling and evaluation

For sampling, we used an adapted form of the “stochastic sampler” from Karras et al. [42], which integrates the ODE while optionally injecting noise at each step. Informally, this is done by first stepping to a higher noise level by adding some noise, and then taking a larger denoising step to remove the added noise in addition to the original step size. We adapted the algorithm slightly: we do not compute the second order correction; we added a step scale which scales the gradient; we do not cap the gamma scale which determines the scale of the noise increase. We describe a subset of experiments we performed to optimize the sampling hyperparameters in Supp. Table S1; the most relevant hyperparameters to tune were the number of steps, the amount of churn (“s_churn”), and the step scale.

#### Algorithm 2

Denoising step (adapted from “StochasticSampler” [42])

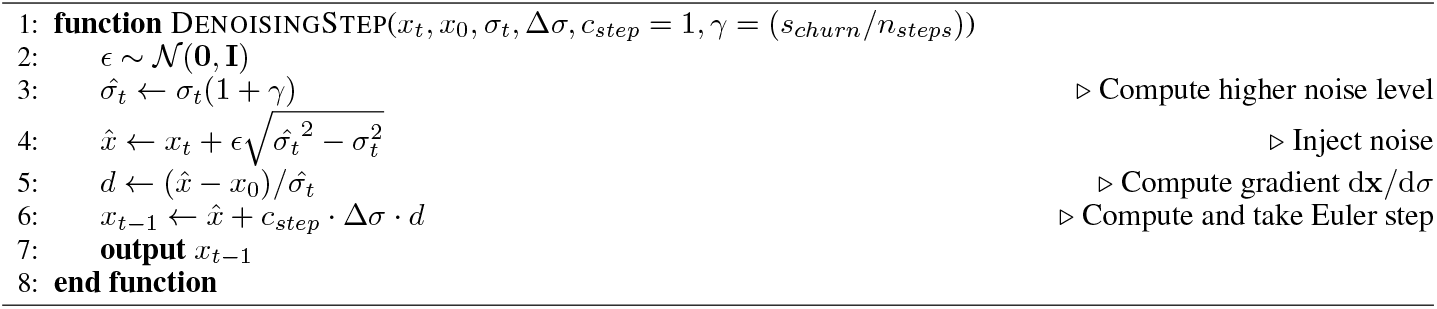

When running the miniMPNN sequence co-design, we sample from the categorical distribution over amino acids independently for each residue and disallow sampling Cys, Unk, and mask tokens. We find that applying transformations on the logits such as temperature or top-p truncation has little benefit on prediction quality in this scheme. Skipping the early portion of the sampling process has little effect on the sequence quality since the structures are so noisy, and it also increases sampling speed significantly; by default we skip miniMPNN co-design for the first 60% of sampling trajectory steps. We also found that not resampling the sequence at every step aids in improving sidechain chemical fidelity, and we used a scheme to linearly anneal the resampling rate from 1 to 0 over the portion of the sampling trajectory where we run miniMPNN co-design.

All self-consistency metrics described are computed with ESMFold [50], and ProteinMPNN where needed [47]. Clustering is performed with maxCluster using single-linkage hierarchical clustering and TM-score threshold of 0.6 [63, 64]. Secondary structure is computed with DSSP [51]. Nearest neighbor searches are computed with Foldseek using the aligned TM score; where we had no hits we report an nnTM of zero [52].

### A.4 Supplemental Figures

**Figure S1.**
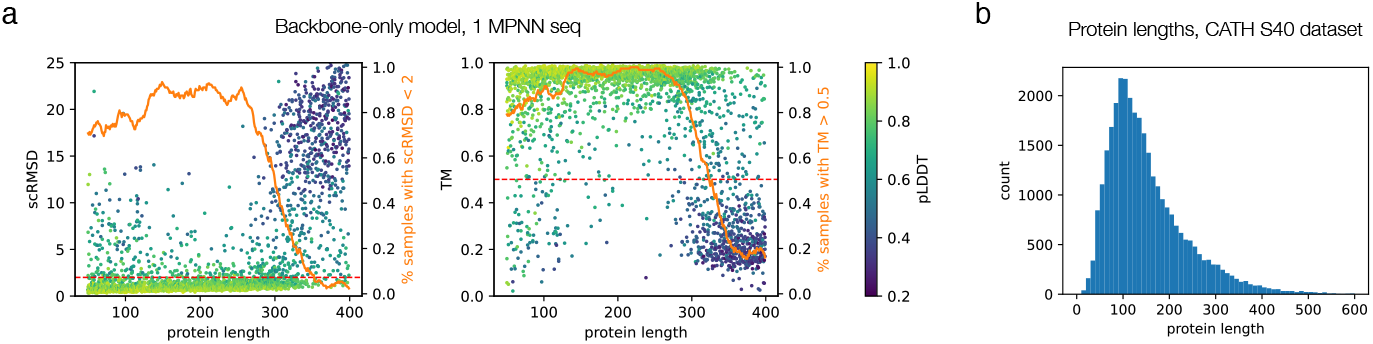
Additional metrics. **(a)** Self-consistency metrics for backbone samples as in Fig. 1, but with only one ProteinMPNN-designed sequence per backbone. **(b)** Distribution of protein lengths in the dataset (excluding proteins with length > 600).

**Figure S2.**
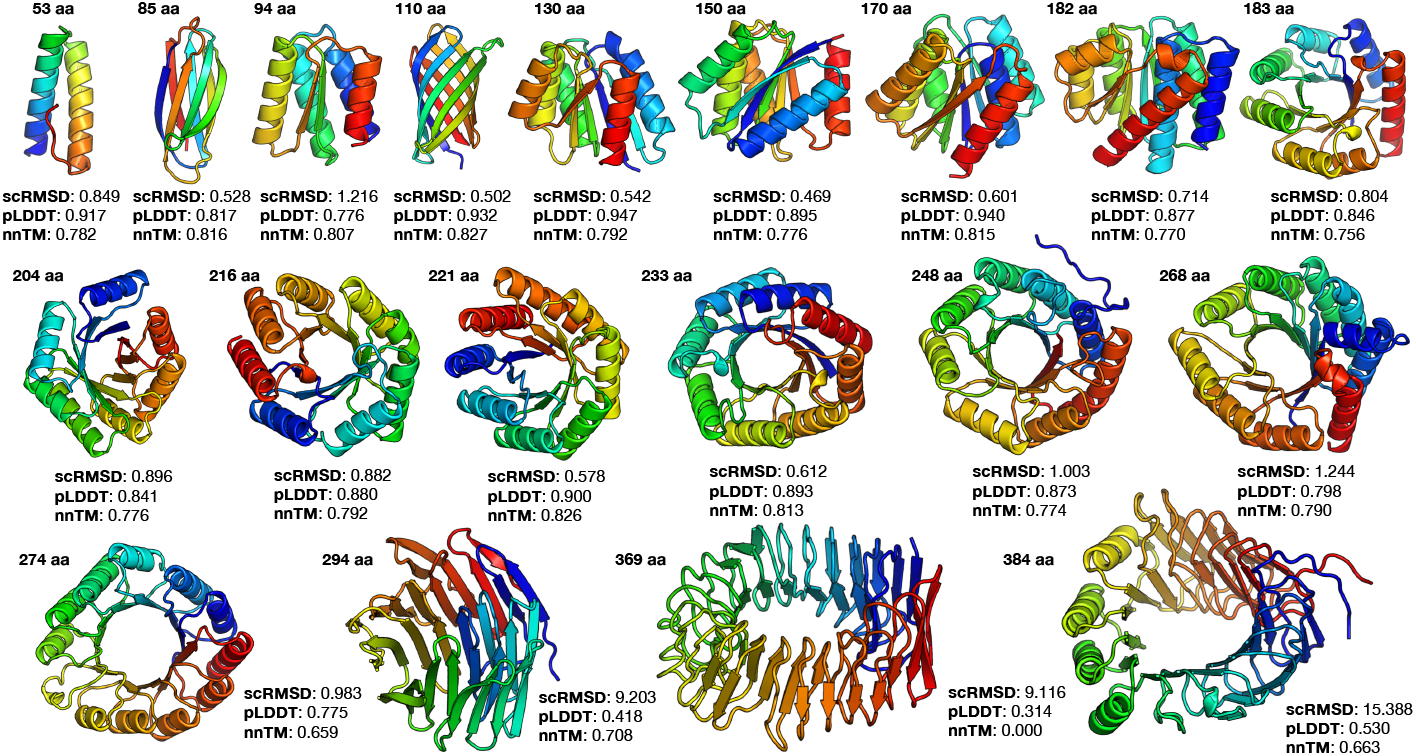
Samples from backbone Protpardelle. Non-cherry-picked raw samples from the backbone model. scRMSD is the best out of 8 ProteinMPNN sequences with ESMFold, with the corresponding pLDDT. Where nnTM is zero, it means we did not find any matches with FoldSeek.

**Figure S3.**
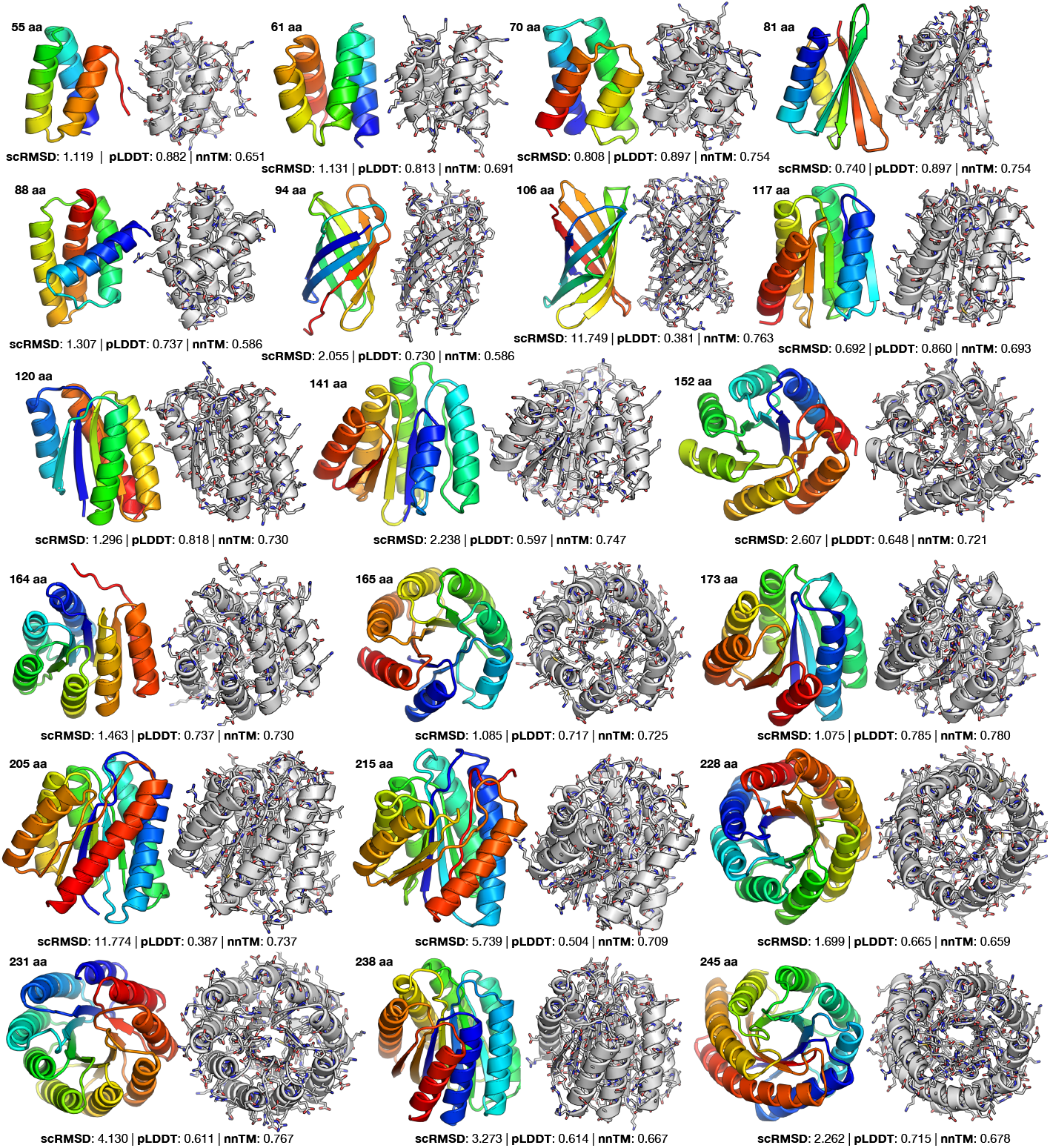
Samples from all-atom Protpardelle. Non-cherry-picked raw samples from the all-atom model. scRMSD is the best out of 8 ProteinMPNN sequences with ESMFold, with the corresponding pLDDT. The structure without sidechains is shown in color, and the same structure with sidechains shown adjacent without color.

**Figure S4.**
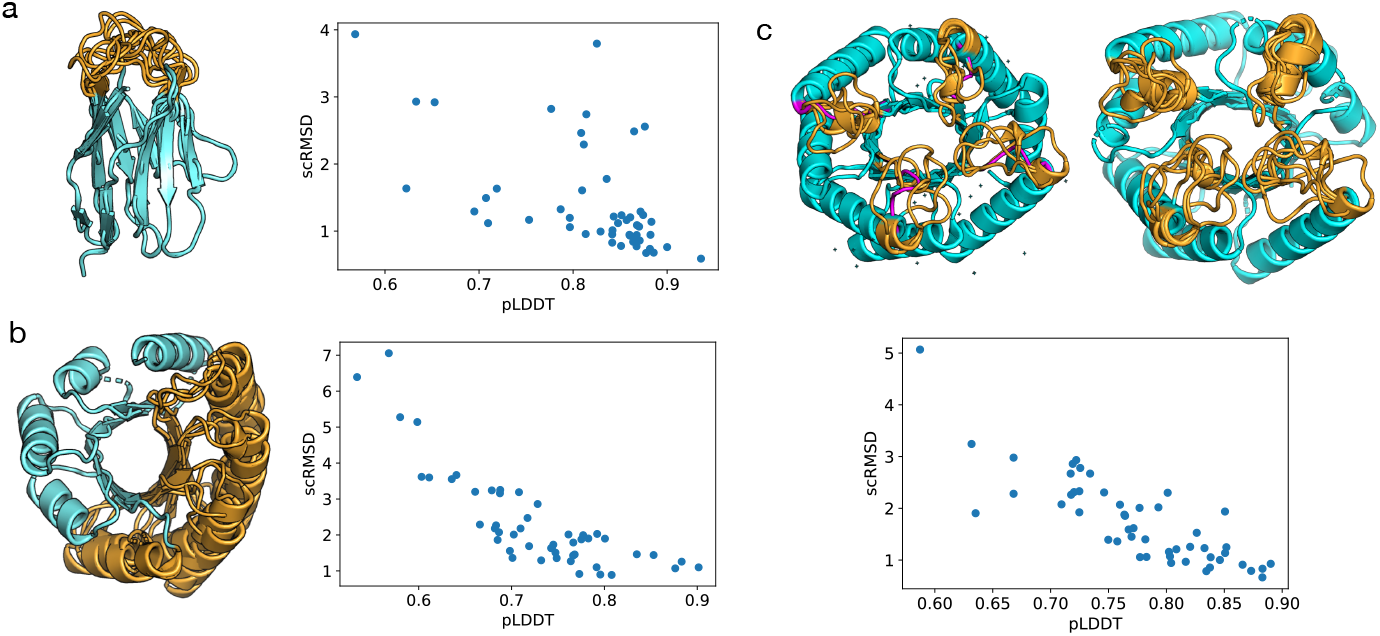
Inpainting ensembles. Inpainting ensembles for **(a)** a loop of a monobody (PDB: 5X2O, chain L) **(b)** half of a *de novo* TIM barrel (PDB: 5BVL) and **(c)** four loops of a *de novo* ovoid TIM barrel (PDB: 7UEK). The conditioning portion is shown in blue, the model generated portion in brown, and the magenta is the original loop. The inpainted structure in (a) and (b) demonstrate same-length inpainted structure, while the length of the loops in (c) are each four residues longer than the original loop length. Sequences were designed with ProteinMPNN and predicted with ESMFold; design success is shown in the plots.

**Table 1:** Experiments on sampling hyperparameters for the all-atom model. For all all-atom results in this paper, we use the “lemon-valley-40, ep2650” model described here. “sampling” describes how many proteins were sampled for each length; the lengths are described in slice format, i.e. 2 for 50:128:1 indicates that 2 proteins at each length from 50 to 128 were sampled. “runtime” describes the number of seconds used to sample all proteins in this set (the number of proteins generated is given in “count”). “steps” describes the number of ODE discretization steps. “schurn” describes the amount of noise that was injected during sampling. “step scale” describes the scale applied to the score before multiplying by the step size. “scRMSD” is the mean and std of scRMSD for all samples in this experiment. “count” describes the total number of proteins sampled. “bond metric” is the bond length RMSE over all atoms.

| model | sampling | runtime (s) | steps | schurn | step scale | scRMSD ($\mu \pm \sigma$) | count | bond metric |
| --- | --- | --- | --- | --- | --- | --- | --- | --- |
| amber-grass-27, ep1900 | 2 for 50:128:1 | 1371.13 | 1000 | 200 | 1 | $9.70 \pm 10.82$ | 156 | 0.11 |
| amber-grass-27, ep1900 | 2 for 50:128:1 | 1336.85 | 1000 | 200 | 1.2 | $6.19 \pm 8.43$ | 156 | 0.09 |
| amber-grass-27, ep1900 | 2 for 50:128:1 | 1341.57 | 1000 | 200 | 1.5 | $5.66 \pm 7.57$ | 156 | 0.08 |
| amber-grass-27, ep1900 | 2 for 50:128:1 | 1617.17 | 1000 | 400 | 1 | $8.00 \pm 9.91$ | 156 | 0.11 |
| amber-grass-27, ep1900 | 2 for 50:128:1 | 1359.52 | 1000 | 400 | 1.2 | $4.64 \pm 6.62$ | 156 | 0.08 |
| amber-grass-27, ep1900 | 2 for 50:128:1 | 1353.33 | 1000 | 400 | 1.5 | $4.27 \pm 5.74$ | 156 | 0.07 |
| amber-grass-27, ep1900 | 2 for 50:128:1 | 1857.78 | 1500 | 400 | 1.2 | $4.83 \pm 7.13$ | 156 | 0.08 |
| amber-grass-27, ep1900 | 2 for 50:128:1 | 2418.27 | 2000 | 400 | 1.2 | $8.81 \pm 11.33$ | 156 | 0.20 |
| amber-grass-27, ep1900 | 2 for 50:128:1 | 1003.19 | 500 | 200 | 1.2 | $3.20 \pm 4.57$ | 156 | 0.08 |
| amber-grass-27, ep1900 | 2 for 50:256:1 | 3683.36 | 1000 | 100 | 1 | $23.57 \pm 17.56$ | 412 | 0.16 |
| amber-grass-27, ep1900 | 2 for 50:256:1 | 3609.14 | 1000 | 100 | 1.2 | $20.98 \pm 18.66$ | 412 | 0.12 |
| amber-grass-27, ep1900 | 2 for 50:256:1 | 3538.07 | 1000 | 100 | 1.5 | $15.98 \pm 18.13$ | 412 | 0.09 |
| amber-grass-27, ep1900 | 2 for 50:256:1 | 1209.80 | 200 | 0 | 1 | $15.59 \pm 11.51$ | 412 | 0.16 |
| amber-grass-27, ep1900 | 2 for 50:256:1 | 1058.20 | 200 | 0 | 1.2 | $11.93 \pm 14.31$ | 412 | 0.12 |
| amber-grass-27, ep1900 | 2 for 50:256:1 | 1093.13 | 200 | 0 | 1.5 | $16.43 \pm 12.69$ | 412 | 0.15 |
| amber-grass-27, ep1900 | 2 for 50:256:1 | 1305.67 | 200 | 100 | 1 | $14.56 \pm 13.73$ | 412 | 0.17 |
| amber-grass-27, ep1900 | 2 for 50:256:1 | 1256.96 | 200 | 100 | 1.2 | $10.05 \pm 13.69$ | 412 | 0.17 |
| amber-grass-27, ep1900 | 2 for 50:256:1 | 1224.22 | 200 | 100 | 1.5 | $23.32 \pm 13.51$ | 412 | 0.31 |
| amber-grass-27, ep1900 | 2 for 50:256:1 | 1266.96 | 200 | 200 | 1 | $14.93 \pm 13.39$ | 412 | 0.16 |
| amber-grass-27, ep1900 | 2 for 50:256:1 | 1232.18 | 200 | 200 | 1.2 | $10.11 \pm 13.76$ | 412 | 0.17 |
| amber-grass-27, ep1900 | 2 for 50:256:1 | 1237.49 | 200 | 200 | 1.5 | $22.75 \pm 13.31$ | 412 | 0.33 |
| amber-grass-27, ep1900 | 2 for 50:256:1 | 1326.97 | 200 | 50 | 1 | None | None | 0.17 |
| amber-grass-27, ep1900 | 2 for 50:256:1 | 1255.39 | 200 | 50 | 1.2 | $10.58 \pm 13.61$ | 412 | 0.15 |
| amber-grass-27, ep1900 | 2 for 50:256:1 | 1288.46 | 200 | 50 | 1.5 | $19.16 \pm 14.53$ | 412 | 0.28 |
| amber-grass-27, ep1900 | 2 for 50:256:1 | 2046.42 | 500 | 0 | 1 | $16.73 \pm 13.99$ | 412 | 0.17 |
| amber-grass-27, ep1900 | 2 for 50:256:1 | 2004.81 | 500 | 0 | 1.2 | $12.45 \pm 15.69$ | 412 | 0.10 |
| amber-grass-27, ep1900 | 2 for 50:256:1 | 1971.94 | 500 | 0 | 1.5 | $15.42 \pm 15.09$ | 412 | 0.10 |
| amber-grass-27, ep1900 | 2 for 50:256:1 | 2266.17 | 500 | 100 | 1 | $18.28 \pm 16.92$ | 412 | 0.15 |
| amber-grass-27, ep1900 | 2 for 50:256:1 | 2117.65 | 500 | 100 | 1.2 | $14.93 \pm 18.06$ | 412 | 0.19 |
| amber-grass-27, ep1900 | 2 for 50:256:1 | 2171.80 | 500 | 100 | 1.5 | $13.93 \pm 18.20$ | 412 | 0.09 |
| amber-grass-27, ep1900 | 2 for 50:256:1 | 2136.91 | 500 | 200 | 1 | $17.86 \pm 16.92$ | 412 | 0.15 |
| amber-grass-27, ep1900 | 2 for 50:256:1 | 2147.56 | 500 | 200 | 1.2 | $12.47 \pm 17.54$ | 412 | 0.19 |
| amber-grass-27, ep1900 | 2 for 50:256:1 | 2115.05 | 500 | 200 | 1.5 | $11.42 \pm 16.11$ | 412 | 0.09 |
| icy-glade-31, ep850 | 4 for 50:256:1 | 2822.46 | 500 | 200 | 1.2 | $5.46 \pm 5.45$ | 824 | 0.12 |
| lemon-shape-51, ep1200 | 8 for 50:256:1 | 3683.67 | 500 | 200 | 1.2 | $4.47 \pm 3.78$ | 1300 | 0.11 |
| **lemon-valley-40, ep2650** | 8 for 50:256:1 | 3523.56 | 500 | 200 | 1.2 | $3.46 \pm 4.33$ | 1648 | 0.11 |
| rare-valley-39, ep1400 | 16 for 50:256:1 | 7774.16 | 500 | 200 | 1.2 | $4.85 \pm 4.16$ | 3296 | 0.11 |
| rich-tree-35, ep1100 | 4 for 50:256:1 | 3161.89 | 500 | 200 | 1.2 | $5.00 \pm 4.22$ | 824 | 0.11 |
| rich-tree-35, ep1300 | 4 for 50:256:1 | 2764.66 | 500 | 200 | 1.2 | $4.26 \pm 3.56$ | 824 | 0.11 |

